# OrthoFiller: utilising data from multiple species to improve the completeness of genome annotations

**DOI:** 10.1101/098566

**Authors:** Michael P. Dunne, Steven Kelly

## Abstract

**Backround:** Complete and accurate annotation of sequenced genomes is of paramount importance to their utility and analysis. Differences in gene prediction pipelines mean that genome sequences for a species can differ considerably in the quality and quantity of their predicted genes. Furthermore, genes that are present in genome sequences sometimes fail to be detected by computational gene prediction methods. Erroneously unannotated genes can lead to oversights and inaccurate assertions in biological investigations, especially for smaller-scale genome projects which rely heavily on computational prediction.

**Results:** Here we present OrthoFiller, a tool designed to address the problem of finding and adding such missing genes to genome annotations. OrthoFiller leverages information from multiple related species to identify those genes whose existence can be verified through comparison with known gene families, but which have not been predicted. By simulating missing gene annotations in real sequence datasets from both plants and fungi we demonstrate the accuracy and utility of OrthoFiller for finding missing genes and improving genome annotation. Furthermore, we show that applying OrthoFiller to existing “complete” genome annotations can identify and correct substantial numbers of erroneously missing genes in these two sets of species.

**Conclusions:** We show that significant improvements in the completeness of genome annotations can be made by leveraging information from multiple species.

## Introduction

Genome sequences have become fundamental to many aspects of biological research. They provide the basis for our understanding of the biological properties of organisms, and enable extrapolation and comparison of information between species. Owing to the increasing availability and affordability [1][2] of whole-genome sequencing technology, genomic data sets are now produced at a rate at which it is infeasible to rely entirely on careful manual curation to annotate a new genome; rather it is taken as given that a considerable portion of the process must be automated.

There has been substantial methodology development in the area of automated gene prediction, with the production of several effective algorithms for identifying genes in *de novo* sequenced genomes [3]. In general, these methods predict genes by learning species-specific characteristics from training sets of manually curated genes. These characteristics include the distribution of intron and exon lengths, intron GC content, exon GC content, codon bias, and motifs associated with the starts and ends of exons (splice donor and acceptor sites, poly-pyrimidine tracts and other features). These characteristics are then used to identify novel genes in raw nucleotide sequences. These prediction methods vary in their performance, as demonstrated by considerable disagreement in the genes and gene models that they predict [4][3]. For example, one study [4] comparing Augustus, GENSCAN, Fgenesh and MAKER, looked at the number of genes predicted on a sample set of *D. melanogaster* assemblies with varying numbers of scaffolds. At the extreme end, with 707 scaffolds, the most frugal prediction (MAKER, with 12687 predicted genes) was almost doubled by the most generous prediction (GENSCAN, with 22679 predicted genes). Thus it is to be expected that genome annotations generated by different research groups using different methodologies will differ considerably in the complement of genes that they contain.

Absent or inaccurate gene models can not only contribute to oversights in biological investigations, they can also lead to false assertions in large-scale genome and cross-species analyses [5]. For example, incorrectly missing gene annotations can be mistakenly interpreted as gene loss, and such interpretations can lead to mistaken inferences about the biological or metabolic properties of an organism. Similarly, missing gene models can lead to errors in gene expression analyses that map and quantify RNA-seq reads using predicted gene models. Here, reads derived from erroneously missing genes, as they have no reference to map to, have the potential to map to the wrong gene leading to errors in transcript abundance estimation.

Much of the cost and effort involved in *de novo* genome annotation can be reduced by leveraging data from other taxa. Moreover, data from disparate taxa have the potential to be used to simultaneously improve a cohort of genome annotations in a mutualistic framework. A number of approaches have been developed to utilise data from other species to improve or assist the process of genome annotation. For example, an automated alignment-based fungal gene prediction (ABFGP) method [6] has been developed for fungal genomes. While this method works well on fungal genomes, it cannot be applied to other taxa and thus has limited general utility.

OrthoFiller aims to simultaneously leverage data from multiple species to mutually improve the genome annotations of all species under consideration. It is designed specifically to find “missing” genes in sets of predicted genes from multiple species. That is, to identify those genes that should be present in a genome's annotation, whose existence can be verified through comparison with known gene families. A standalone implementation of the algorithm is available under the GPLv3 licence at https://github.com/mpdunne/orthofiller.

## Results

### Problem definition, algorithm overview and evaluation criteria

OrthoFiller aims to find genes that are present in a species’ genome, but which have no predicted gene model in the genome annotation for that species. It takes a probabilistic, orthogroup-based approach to gene identification, leveraging information from multiple species simultaneously to improve the completeness of the genome annotations for all species under consideration. OrthoFiller is not designed for *ab initio* gene prediction and requires that each genome under consideration possesses a basic level of annotation, taken to be at least 100 annotated genes. The genomes should ideally be from a set of related species from the same taxonomic group (genus, family, order or class).

A workflow for OrthoFiller is shown in Figure 1. The basic input for the algorithm is a set of genome annotation files in general transfer format (GTF) and a set of corresponding genome sequence files in FASTA format. Protein sequences are extracted from the genome FASTA files using the coordinates in the GTF files and a user-selected translation table. The predicted proteomes from the submitted species are clustered into orthogroups using OrthoFinder [7], the protein sequences of each orthogroup are aligned and the source nucleotide sequences for these proteins are threaded back through the protein multiple sequence alignment to create multiple sequence alignments of the nucleotide sequences of each orthogroup. Each nucleotide alignment is used to build a hidden Markov model (HMM) that is used to search the complete genome sequence of each species under consideration. The scores of these HMMs are used to learn the score distributions of true positive and false positive HMM hits (see methods). Each hit to an HMM that does not overlap with an existing predicted gene is subject to filtration using species-specific parameters that have been learned for true and false positive hits. Each hit that survives this filtration is considered to be a potential genic region, or *hint*. The algorithm then attempts to build gene models around these hints, using the Augustus [8] gene finder. Gene models constructed by Augustus are subject to two successive rounds of assessment and filtration. Firstly, the predicted gene models are compared against the hints that were used to inform them: if the gene model and its source hint are not sufficiently similar (see methods), the gene model is considered to be unrelated to the hint, and thus to the orthogroup used to inform its prediction. Secondly, the newly predicted genes that satisfy the first criterion are subject to orthogroup inference using the full set of existing and newly predicted genes. Those newly predicted genes that are clustered in an orthogroup whose HMM was used to predict them are then accepted as *bona fide* genes and added to the genome annotation. Thus genes predicted by OrthoFiller satisfy stringent orthology based criteria for inclusion.

**Figure 1:**
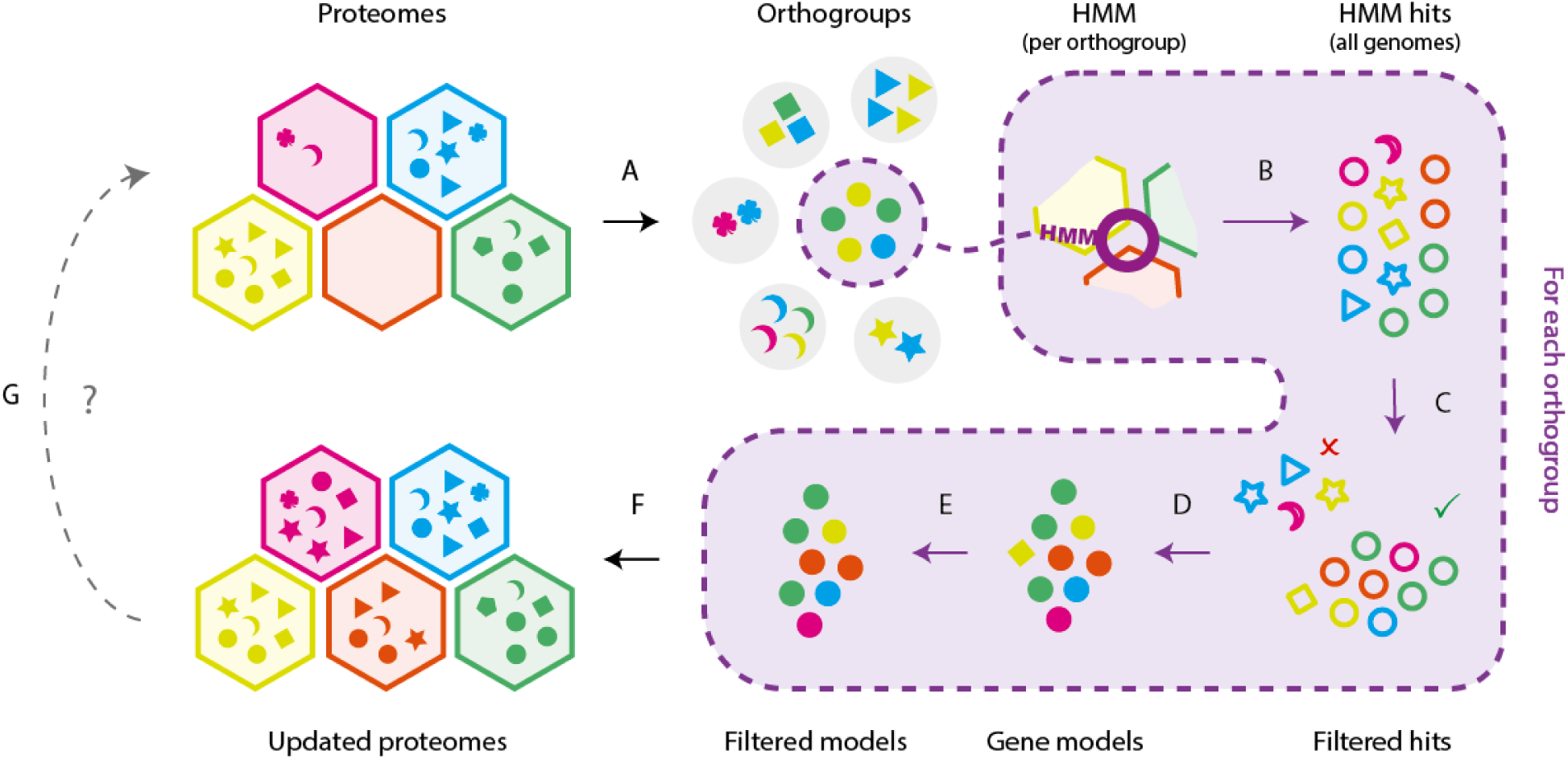
Workflow diagram for the OrthoFiller algorithm. **A)** Proteomes are subdivided into orthogroups using OrthoFinder. **B)** Protein sequences in each orthogroup are subject to multiple sequence alignment, back-translated to DNA and used to create hidden Markov models (HMMs). These HMMs used to search each genome in the set. **C)** The set of hits are evaluated and filtered to remove low quality hits. **D)** Gene models are constructed around each retained hit using Augustus. **E)** The new gene models are compared to the hints that were used to generate them, and filtered to remove those which bear in sufficient similarity to the hints. **F)** The filtered genes are clustered into orthogroups and genes that are successfully placed into the orthogroup that was used to identify them are retained. **G)** The process may be run once, or iteratively until no further genes are found.

To demonstrate the utility of OrthoFiller on real data it was applied independently to two sets of species. Set A comprised five fungal genomes (Table 1) and Set B comprised five plant genomes (Table 2), sourced from the Joint Genome Institute (JGI) and the Saccharomyces Genome Database (SGD) [9][10][11][12]. OrthoFiller was assessed using these datasets in two ways: first via simulating an incomplete genome annotation by randomly removing entries from the genome annotation of one species from each set, and assessing the accuracy of OrthoFiller in recovering the removed genes; second by application of OrthoFiller to the complete datasets and validating the novel detected genes through analysis of publicly available RNA-seq data.

**Table 1:**
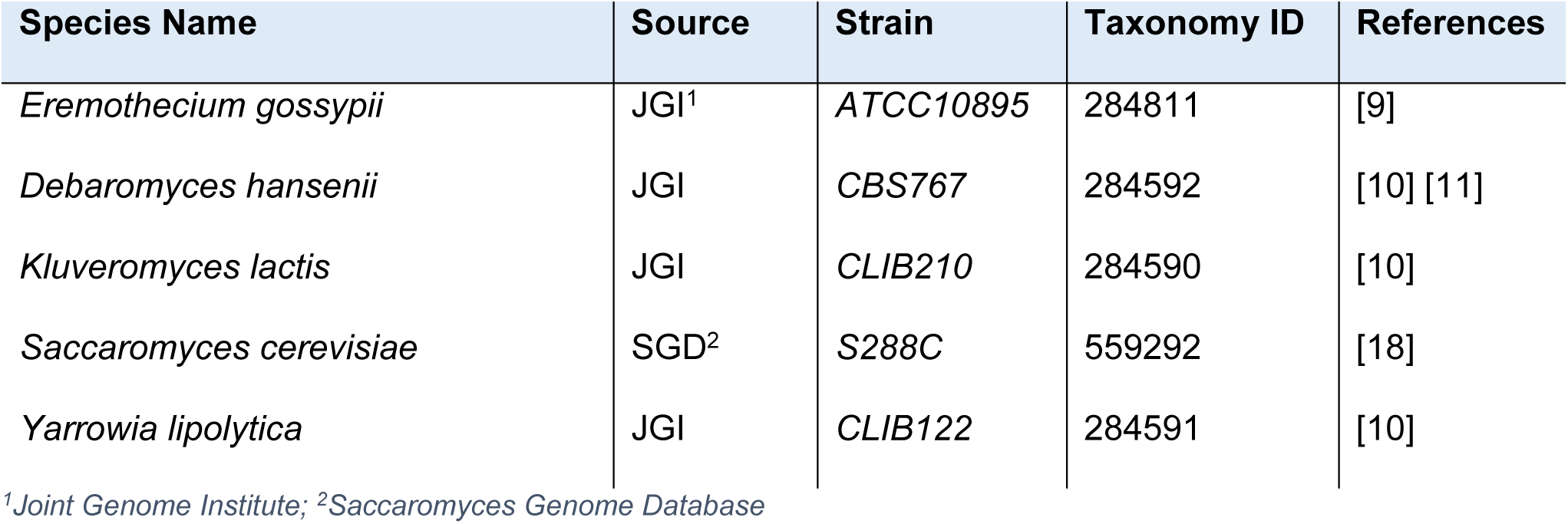
Species Set A, fungal species used for algorithm validation.

| Species Name | Source | Strain | Taxonomy ID | References |
| --- | --- | --- | --- | --- |
| *Eremothecium gossypii* | JGI<sup>1</sup> | ATCC10895 | 284811 | [9] |
| *Debaromyces hansenii* | JGI | CBS767 | 284592 | [10] [11] |
| *Kluveromyces lactis* | JGI | CLIB210 | 284590 | [10] |
| *Saccaromyces cerevisiae* | SGD<sup>2</sup> | S288C | 559292 | [18] |
| *Yarrowia lipolytica* | JGI | CLIB122 | 284591 | [10] |

<sup>1</sup>Joint Genome Institute; <sup>2</sup>Saccaromyces Genome Database

**Table 2:** Species Set B, plant species used for algorithm validation.

| Species Name | Source | Version | Taxonomy ID | References |
| --- | --- | --- | --- | --- |
| *Arabidopsis thaliana* | JGI | TAIR10 | 3702 | [12] |
| *Brassica rapa* | JGI | v1.3 | 3711 | [12] |
| *Carica papaya* | JGI | ASGPBv0.4 | 3649 | [12] |
| *Capsella rubella* | JGI | V1.0 | 81985 | [12] |
| *Theobroma cacao* | JGI | V1.1 | 3641 | [12] |

Two measures were used to assess the quality of recovered genes: the protein F-score and the orthogroup F-score, both defined in the methods section. These scores were calculated for all genes identified by OrthoFiller, by comparing the recovered gene with the removed gene and assuming that the original removed gene model was correct. Genes that are unique to the test species that lack homologues in other species were not analysed in this test, as OrthoFiller was designed to find evolutionarily conserved genes. As there were no publicly-available comparable methods that perform the same task as OrthoFiller, the method was assessed in comparison to performing the analysis without conducting the OrthoFiller evaluation and filtration steps. i.e. accepting all identified gene models that did not overlap an existing gene.

### Evaluation of OrthoFiller on *S. cerevisiae* after removal of 10% of gene annotations

Figure 2 and Table 3 show the results of running OrthoFiller on the set of fungal species shown in Table 1 after random removal of 10% of “discoverable” genes (genes that were contained in an orthogroup with at least one gene from another species) from the predicted complement of genes in S. *cerevisiae* (i.e. 528 nuclear encoded gene annotations were deleted from a total set of 5288 discoverable genes).

**Figure 2:**
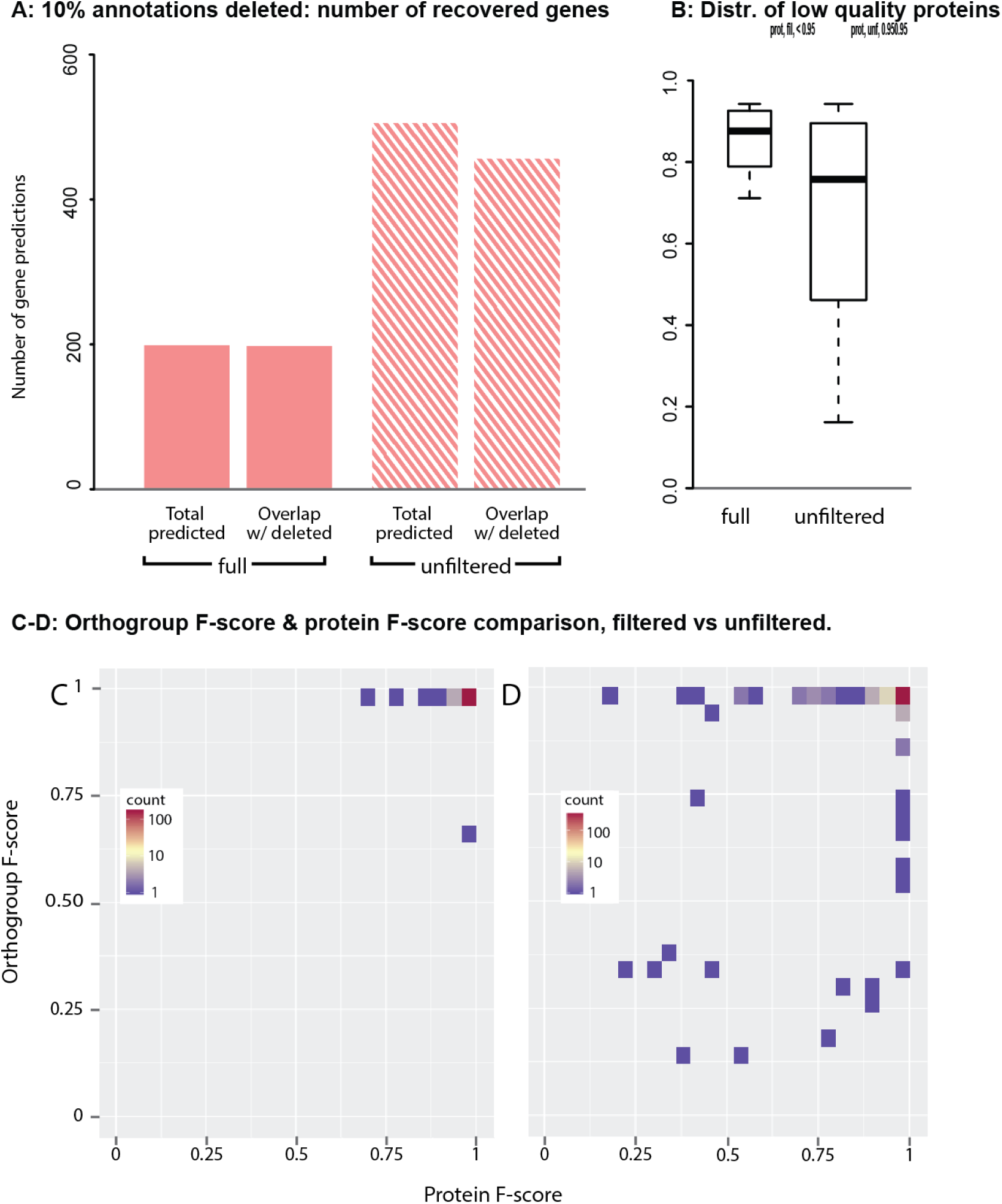
Performance of OrthoFiller on *S. cerevisiae* genome with 10% of annotated genes removed. **A)** Using OrthoFiller 197 genes were found whose genomic locations matched any of the 528 deleted genes. In the absence of OrthoFiller filtration this increased to 447 genes identified that overlap any part of a deleted gene. **B**) A boxplot of protein F-scores for genes predicted using OrthoFiller, or in the absence of OrthoFiller filtration, that had a protein F-score of ≤0.95. **C**) Density plot showing the protein and orthogroup F-scores for all recovered genes using OrthoFiller. **D**) Density plot showing the protein and orthogroup F-scores for all recovered genes in the absence of OrthoFiller filtration.

**Table 3:** Recovery of removed genes in *S. cerevisiae*.

|  | 10% annotations removed |  | 90% annotations removed |  |
| --- | --- | --- | --- | --- |
|  | OrthoFiller | Unfiltered | OrthoFiller | Unfiltered |
| No. genes removed | 528 | 528 | 4759 | 4759 |
| Total genes found | 197 | 503 | 1529 | 4325 |
| Found genes which overlap removed genes | 196 | 447 | 1529 | 4156 |
| Total recovered genes | 196 | 440 | 1528 | 4116 |
| Number of split genes | 0 | 7 | 1 | 34 |
| Mean pF score of found genes | 0.99 | 0.97 | 0.99 | 0.97 |
| Mean oF score of found genes | 1.00 | 0.98 | 0.98 | 0.96 |
| High-quality found genes (pF≥0.95) | 190 | 411 | 1455 | 3801 |
| Lower-quality found genes (pF<0.95) | 6 | 36 | 74 | 355 |
| Mean pF-score of lower-quality genes | 0.85 | 0.68 | 0.87 | 0.69 |
| % of lower-quality genes with oF≥0.95 | 100.0% | 69.4% | 91.9% | 68.2% |

After running OrthoFiller, a total of 197 genes were predicted in the genome of *S. cerevisiae* that were not present in the submitted genome annotation file. Of these, 196 overlapped with genes that were deleted from the original annotation and one was not present in the original annotation (37.1% of 528, Figure 2A). In total, 190 of the 196 found genes (96.9%) were recovered to high accuracy (protein F-score ≥ 0.95). The mean protein F-score of the remaining 6 genes of lower accuracy (protein F-score < 0.95) was 0.85 (Figure 2B). All of the genes that had lower gene model accuracy were placed in exactly the same orthogroup as expected when the sequences were subjected to orthogroup inference. Thus, although 6 of the gene models differed from the original reference gene model, this difference was not sufficient to disrupt downstream identification of orthologous genes.

To provide a comparison, in the absence of the OrthoFiller evaluation steps a total of 503 genes were identified, of which 447 overlapped with genes that were deleted from the original annotation and 56 were not predicted as genes in the original *S. cerevisiae* genome (Figure 2A). The regions comprising these 447 found genes corresponded to 440 deleted genes (84.6% of 528). This discrepancy in gene number is due to genes which were recovered, but whose recovered versions were split into multiple parts. There were 7 such split genes. In total, 411 (91.9% of 447) of the genes that overlapped with genes present in *S. cerevisiae* genome annotation were genes recovered with high accuracy (protein F-score ≥ 0.95) and the mean protein F-score of those recovered to a lower accuracy was 0.68 (Figure 2B), considerably lower than in the filtered case. Of these 36 lower-quality genes, 11 (30.5%) had an orthogroup F-score less than or equal to 0.95. Moreover, 10 of these genes were sufficiently mis-predicted that they failed to be placed in an orthogroup, or were placed in an orthogroup that shared no members with the orthogroup that contained the original gene. Thus in the absence of OrthoFiller filtration, more genes were recovered but 6 genes were fragmented, 10 of the found genes bore insufficient similarity to the reference gene to facilitate orthogroup inference, and 26 were sufficiently mis-predicted that the results of orthogroup inference was altered.

Figures 2C-D show the distribution of orthogroup F-scores versus protein F-scores obtained following application of OrthoFiller to this test dataset. The majority of recovered genes had both high protein and orthogroup F-scores (Figure 2C): 189 out of 196 genes (96.4%) had both F-scores ≥ 0.95. This indicates that the majority of predicted genes are identical (or nearly identical) to the original removed gene and that when subject to orthogroup inference they were clustered in the correct orthogroup. Imperfect protein F-scores can be explained by discrepancies in intron/exon and start/stop codon choices between the removed and recovered gene models. Imperfect orthogroup F-scores were due to fluctuations in orthogroup membership. Figure 2D shows the results in the absence of OrthoFiller processing. In this case, 399 of 447 genes (89.3%) were of dually high quality.

In particular, there were 5 predicted genes with both a low (< 0.5) protein and orthogroup F-score, indicating those predicted genes were sufficiently incorrect to cause errors in orthologous gene identification. Thus, although OrthoFiller does not recover all deleted genes (37% of removed genes), application of OrthoFiller resulted in the recovery of high-quality gene annotations that contain few (in this example there are none) incorrectly predicted genes.

### Evaluation of OrthoFiller on *S. cerevisiae* after removal of 90% of gene annotations

Figure 3 and Table 3 show the performance statistics for OrthoFiller using a version of *S. cerevisiae* genome where 90% of gene annotations were removed. This represents an extreme case where a genome has minimal annotation. The full details of detection of the deleted genes at different stages in the OrthoFiller algorithm are shown in Supplemental Figure 3. Here, application of OrthoFiller resulted in the identification of 1529 genes that overlapped with 1528 of the removed genes (32.1%, Figure 3A). One of the genes was split into two parts. Of the found genes, 1455 (95.1%) were recovered with a protein F-score of 0.95 or greater. Of the 74 genes with lower protein F-scores (Figure 3B), only 6 (8.1%) had an orthogroup F-score < 0.95. As before, although these gene models differed from the original reference gene model, this difference was not sufficient to disrupt downstream identification of orthologous genes.

**Figure 3:**
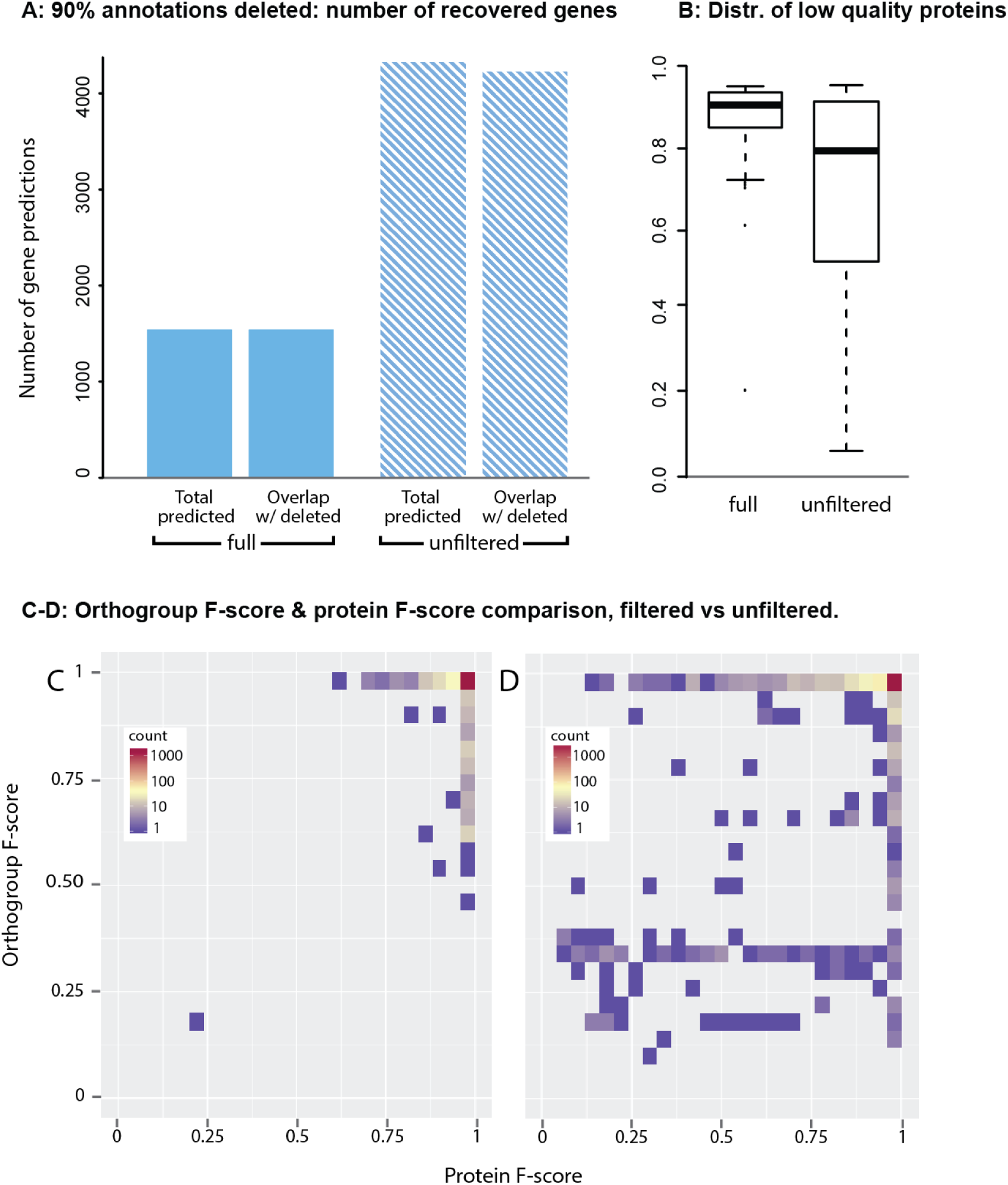
Performance of OrthoFiller on S. *cerevisiae* genome with 90% of annotated genes removed. **A)** Using OrthoFiller 1529 genes were found which overlapped any of the 4759 deleted genes. In the absence of OrthoFiller filtration this increased to 4156 genes. **B)** A boxplot of protein F-scores for genes predicted using OrthoFiller, or in the absence of OrthoFiller filtration, that had a protein F-score of ≤0.95. **C)** Density plot showing the protein and orthogroup F-scores for all recovered genes using OrthoFiller. **D)** Density plot showing the protein and orthogroup F-scores for all recovered genes in the absence of OrthoFiller filtration.

In the absence of OrthoFiller filtration, 4325 genes were found, of which 4116 overlapped the removed genes. Of the removed genes, 4156 were recovered, of which 64 genes were split. 3801 of the found genes had a protein F-score ≥ 0.95 (87.9%). Of the 355 genes with lower protein F-scores, 113 had an orthogroup F-score lower than 0.95, and 97 were sufficiently mis-predicted that they failed to be placed in any orthogroup at all, or in an orthogroup completely different to the one that was used to find them.

Figures 3C-D show the distribution of orthogroup F-scores versus protein F-scores for recovery in the 90% removal case. Figure 3C shows that most genes were recovered well, with 1367 of 1529 (89.4%) genes predicted correctly and placed in the correct orthogroup when subject to orthogroup inference (protein F-score ≥ 0.95, orthogroup F-score ≥ 0.95). Interestingly, there are many genes that are predicted correctly but are placed into a slightly different orthogroup to what was expected. This is due to changes in orthogroup membership caused by the many still-missing genes. Thus, although the input datasets are dramatically different the performance characteristics of OrthoFiller on the 10% and 90% datasets are broadly consistent (e.g. 37.1% and 32.1% recovery respectively, 96.9% and 95.1% high-accuracy recoveries respectively).

### Evaluation of OrthoFiller on *A. thaliana* after removal of 10% of gene annotations

As it could be argued that fungal genomes present an easier challenge, an additional demonstration of the utility of OrthoFiller on an alternative group of organisms was also conducted. Here the analogous test of the method was applied to a set of five land plant genomes (Table 2). Table 4 and Figure 4 show performance statistics from application of OrthoFiller to the *A. thaliana* genome with 10% (3168) gene annotations removed. Out of the 1097 genes that were output by OrthoFiller, 982 overlapped removed genes. A total of 908 of the original genes were recovered, of which 67 were recovered but split into multiple parts (7.4%). Of the found genes, 416 (42.4%) had a protein F-score of 0.95 or higher, and of the lower quality genes, 56.5% had orthogroup F-scores of 0.95 or higher, and 52.5% were placed into exactly the same orthogroup as the one used to predict them. The mean protein F-score of lower-quality genes was 0.60. Thus similar to the fungal dataset, application of OrthoFiller resulted in the identification of 31.0% of the removed genes, with 42.4% being of gene model accuracy (assuming the deleted gene to be true).

**Figure 4:**
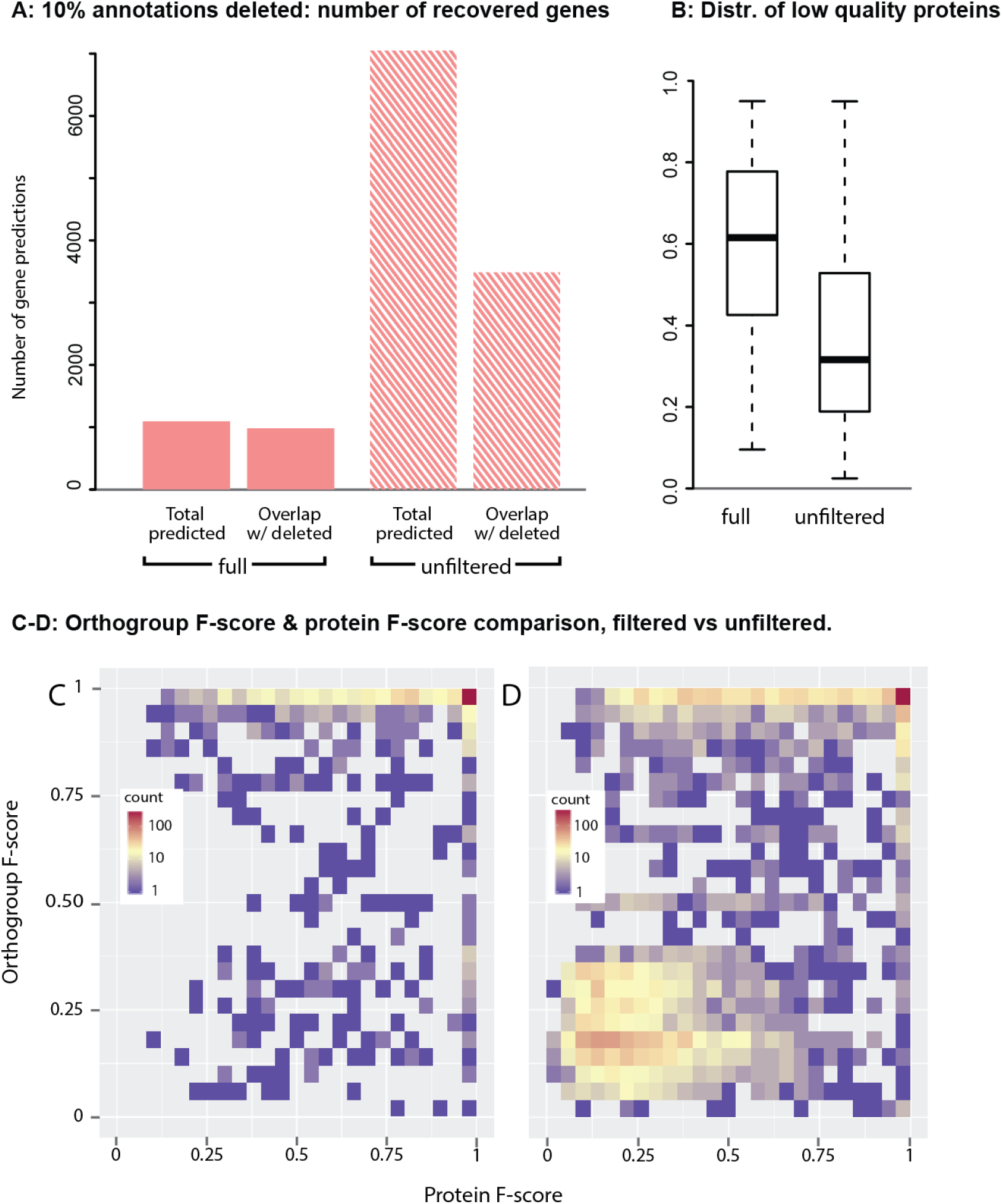
Performance of OrthoFiller on *A. thaliana* genome with 10% of annotated genes removed. **A)** Using OrthoFiller 982 genes were found which overlapped any of the 3168 deleted genes. In the absence of OrthoFiller filtration this increased to 3484 genes. **B)** A boxplot of protein F-scores for genes predicted using OrthoFiller, or in the absence of OrthoFiller filtration, that had a protein F-score of ≤0.95. **C)** Density plot showing the protein and orthogroup F-scores for all recovered genes using OrthoFiller. **D)** Density plot showing the protein and orthogroup F-scores for all recovered genes in the absence of OrthoFiller filtration.

**Table 4:**
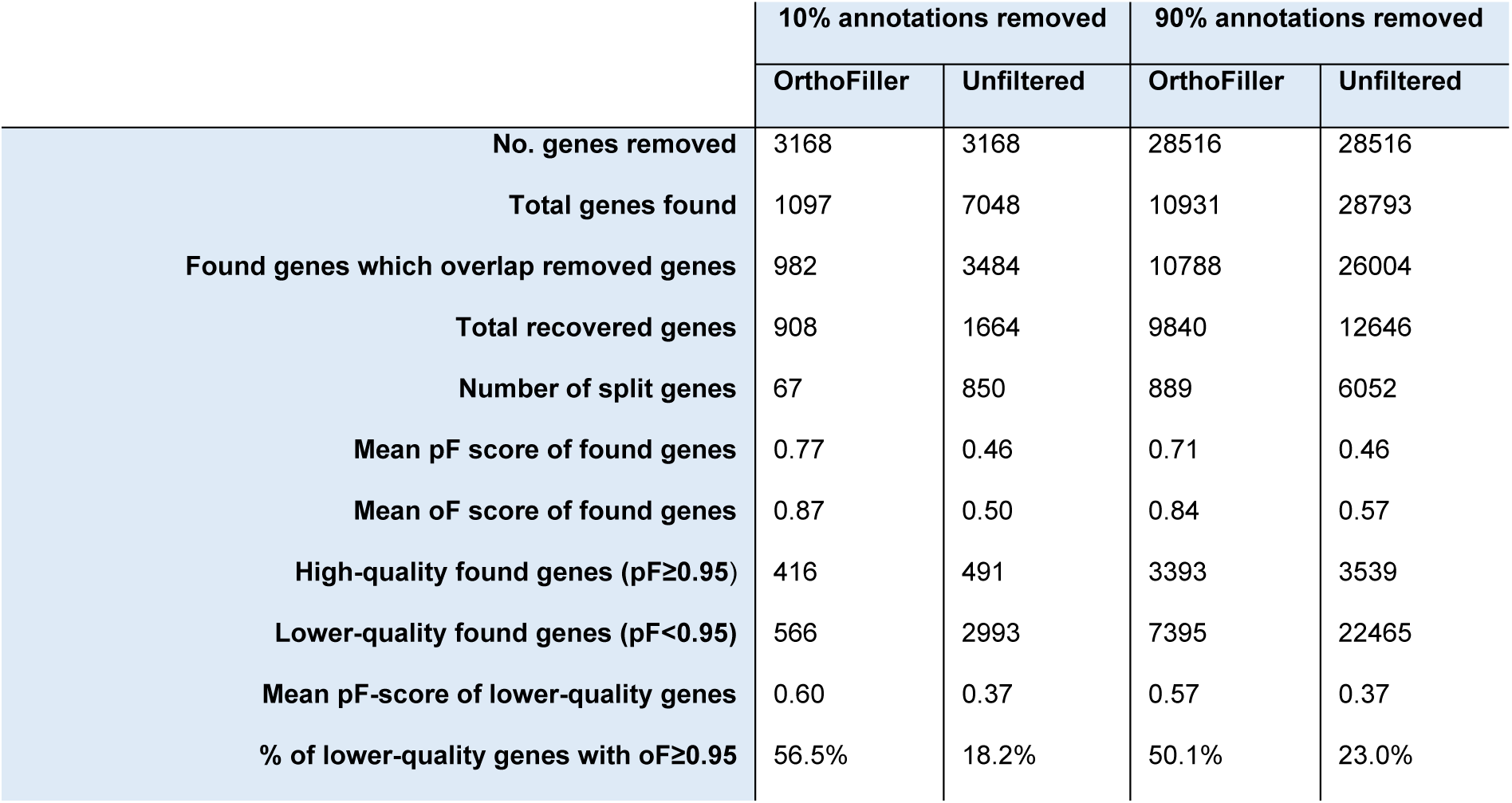
Recovery of removed genes in A. *thaliana*.

|  | 10% annotations removed |  | 90% annotations removed |  |
| --- | --- | --- | --- | --- |
|  | OrthoFiller | Unfiltered | OrthoFiller | Unfiltered |
| No. genes removed | 3168 | 3168 | 28516 | 28516 |
| Total genes found | 1097 | 7048 | 10931 | 28793 |
| Found genes which overlap removed genes | 982 | 3484 | 10788 | 26004 |
| Total recovered genes | 908 | 1664 | 9840 | 12646 |
| Number of split genes | 67 | 850 | 889 | 6052 |
| Mean pF score of found genes | 0.77 | 0.46 | 0.71 | 0.46 |
| Mean oF score of found genes | 0.87 | 0.50 | 0.84 | 0.57 |
| High-quality found genes (pF≥0.95) | 416 | 491 | 3393 | 3539 |
| Lower-quality found genes (pF<0.95) | 566 | 2993 | 7395 | 22465 |
| Mean pF-score of lower-quality genes | 0.60 | 0.37 | 0.57 | 0.37 |
| % of lower-quality genes with oF≥0.95 | 56.5% | 18.2% | 50.1% | 23.0% |

In the absence of OrthoFiller filtration 7048 genes were found, nearly twice as many as were removed. Only 3484 of these overlapped removed genes, of which 491 (14.1%) had a protein F-score of 0.95 or higher. 1664 genes were recovered, of which 850 (51.1%) were split into multiple parts. The mean protein F-score of lower-quality genes was 0.37, and the percentage of lower-quality genes which received an orthogroup F-score of 0.95 or above was 18.2%.

Figures 4C-D show the distribution of orthogroup F-scores versus protein F-scores for recovery in the 10% removal case for *A. thaliana.* Using OrthoFiller, 325 of 902 (33%) of genes had both a very high (≥ 0.95) protein and orthogroup F-score. In the unfiltered case, 324 of the genes had both a high protein and orthogroup F-score, though as a percentage of the total genes found (9.2% of 3484 found genes), the success rate was considerably lower. Conversely, 35 out of 982 (3.6%) had both scores very low (<0.5), compared with 1710 out of 3484 (49.1%) genes in the absence of OrthoFiller filtration. Thus in this case using OrthoFiller considerably reduces the proportion of found genes which are erroneous.

### Evaluation of OrthoFiller on *A. thaliana* after removal of 90% of gene annotations

Performance statistics for the application of OrthoFiller to the 90% depleted *A. thaliana* genome (28516 genes removed) can be seen in Table 4 and Figure 5. Of 10931 found genes, 10788 overlapped removed genes, 3393 of which (31.5%) had protein F-score 0.95 or above. 889 (9.0%) of the recovered genes were split into multiple parts. A total of 9840 out of 28516 (34.5%) removed genes were recovered, though 889 were split into parts (9.0%). Of the lower-quality genes, 50.1% had orthogroup F-score ≥ 0.95, and 46.7% were placed in exactly the right orthogroup. The mean protein F-score of the lower-quality genes was 0.57. Thus having fewer gene models to serve as examples for gene model training resulted in a higher error rate in gene model prediction.

**Figure 5:**
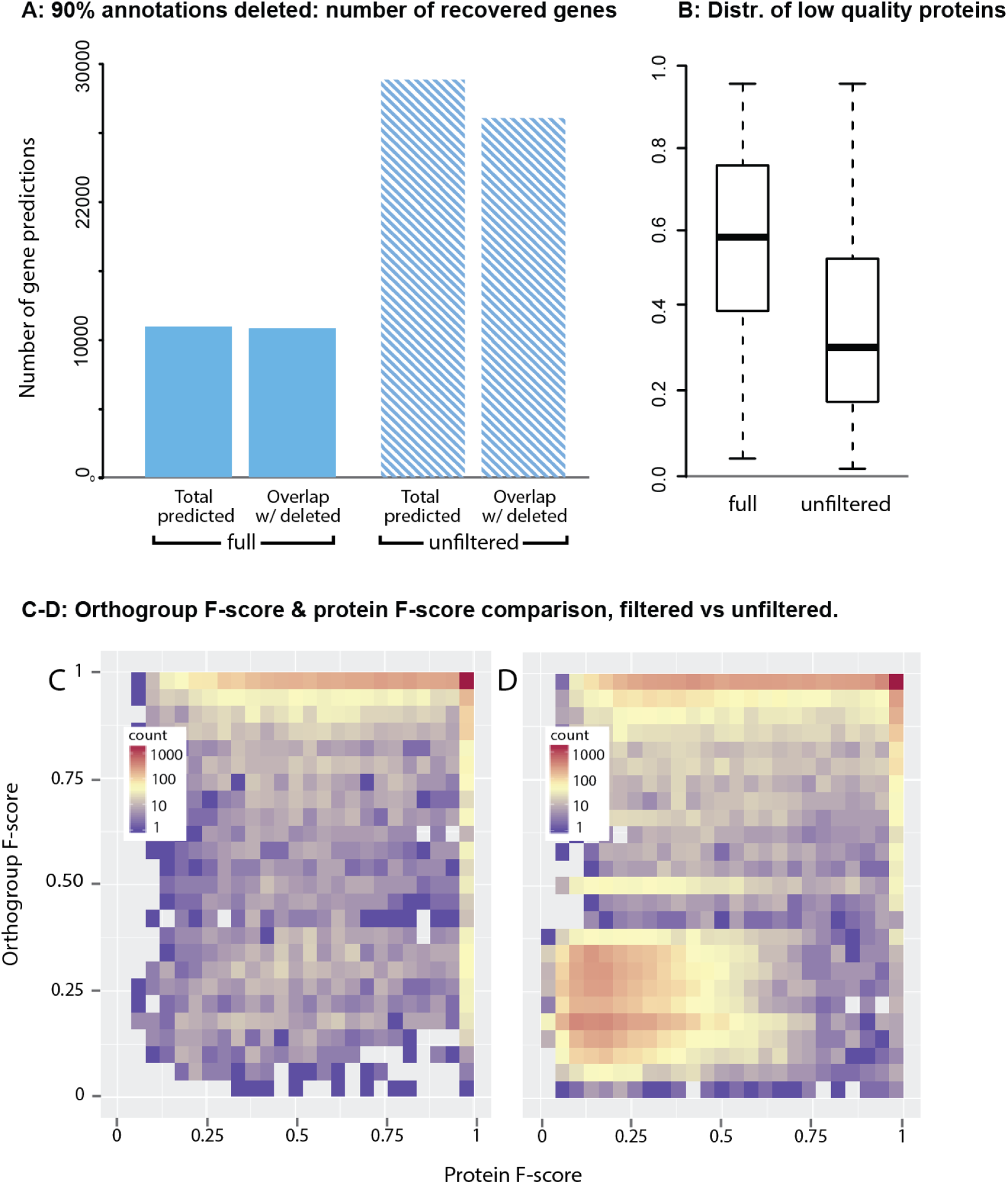
Performance of OrthoFiller on *A. thaliana* genome with 90% of annotated genes removed. **A)** Using OrthoFiller 10788 genes were found which overlapped any of the 28516 deleted genes. In the absence of OrthoFiller filtration this increased to 26204 genes. **B)** A boxplot of protein F-scores for genes predicted using OrthoFiller, or in the absence of OrthoFiller filtration, that had a protein F-score of ≤0.95. **C)** Density plot showing the protein and orthogroup F-scores for all recovered genes using OrthoFiller. **D)** Density plot showing the protein and orthogroup F-scores for all recovered genes in the absence of OrthoFiller filtration.

**Figure 6:**
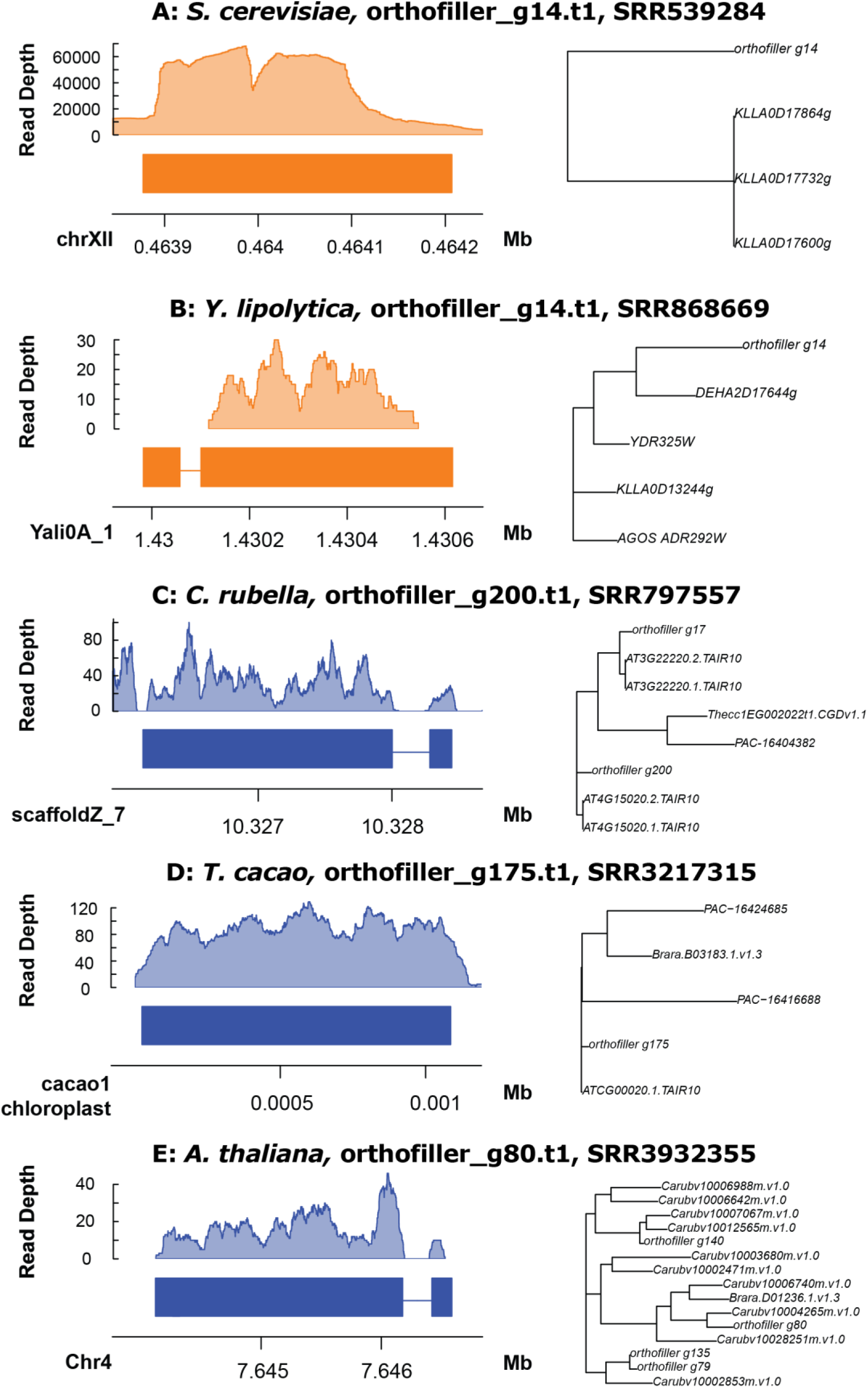
Coverage plots and orthogroup trees for a selection of new genes. Five representative examples of RNAseq coverage on genes predicted using OrthoFiller. Phylogenetic trees demonstrate the relationship of the newly predicted gene to other genes in the orthogroup.

In the absence of OrthoFiller filtration, 28793 genes were predicted, 26004 of which overlapped removed genes. Of these, only 3539 (13.4%) had a protein F-score of 0.95 or above, with just 23% of the lower-quality genes having orthogroup F-score ≥ 0.95. In total 12646 of the 28516 removed genes were recovered, although 6052 of them were split (47.9%). The mean protein F-score of the lower-quality genes was 0.37. This shows that, although slightly more genes were recovered in the unfiltered case, considerably more noise and erroneous predictions are produced.

Figures 5C-D show the distribution of orthogroup F-scores versus protein F-scores for recovery in the 90% removal case for *A. thaliana.* Using OrthoFiller, 2427 of 10788 found genes (22.5%) had both a very high (≥ 0.95) protein and orthogroup F-score, compared with 2413 out of 26004 (9.3%) in the unfiltered case. Conversely, only 5.9% of genes (636 out of 10788) predicted using OrthoFiller had both scores very low (<0.5), compared with 44.7% of genes (11631 out of 26004) in the absence of OrthoFiller filtration. Thus, similarly to with the fungal data set, the performance characteristics of OrthoFiller on the 10% and 90% plant datasets are broadly consistent (e.g. 31.0% and 34.5% recovery respectively, 42.4% and 31.5% high-accuracy recoveries respectively), and both contain a considerably smaller proportion of clearly erroneous genes than would be found without filtering.

### OrthoFiller detects hundreds of conserved genes not present in the reference genome annotations

In addition to testing the ability of OrthoFiller to recover already predicted genes, the algorithm was applied to both of the sets of complete genomes listed in Table 1 and Table 2, to assess the potential for novel genes to be discovered. The number of genes found for each species in each set is listed in Tables 5 and 6. Application of OrthoFiller to the 5 fungal species listed in Table 1 resulted in the detection of 31 novel genes distributed across the 5 species. Further rounds of OrthoFiller gene prediction identified no additional genes to those already found. Application of OrthoFiller to the 5 plant species listed in Table 2 resulted in the identification of 570 individual novel genes in these species.

**Table 5:** Novel genes in fungal species.

| Species Name | Genome size (Mbp) | No. pre-existing genes | No. new genes. |
| --- | --- | --- | --- |
| *E. gossypii* | 9.10 | 4768 | 2 |
| *D. hansenii* | 12.15 | 6272 | 13 |
| *K. lactis* | 10.69 | 5076 | 6 |
| *S. cerevisiae* | 12.16 | 6572 | 2 |
| *Y. lipolytica* | 20.50 | 6447 | 8 |

**Table 6:**
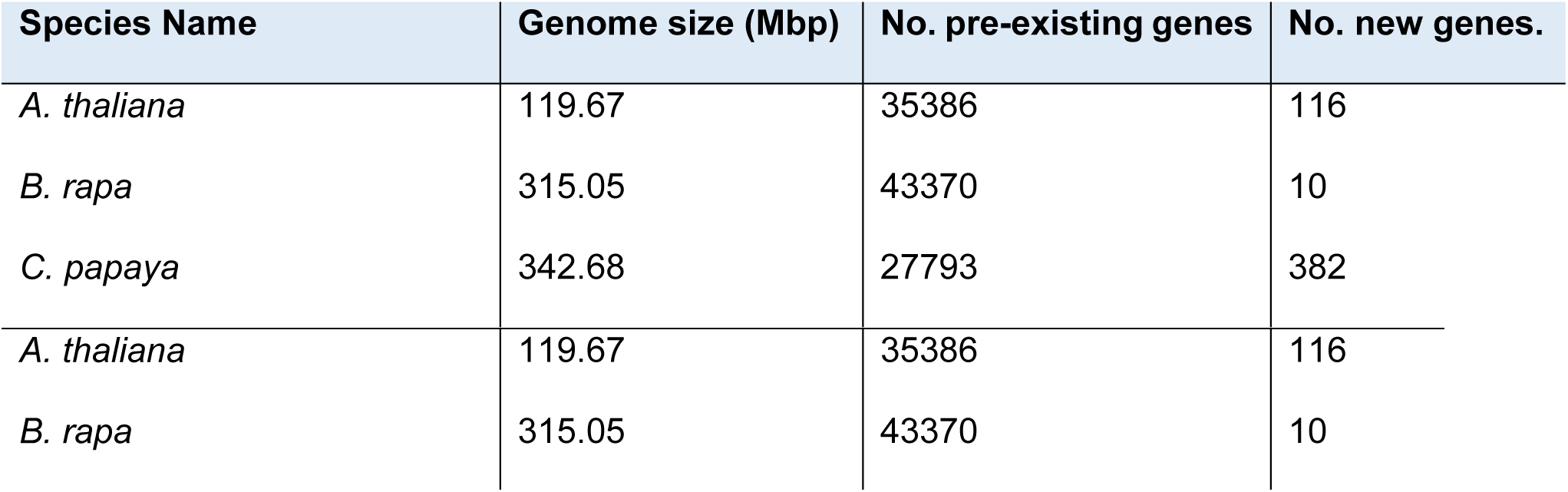
Novel genes in plant species.

To be detected as a novel gene OrthoFiller requires genes to pass rigorous sequence similarity tests to genes in other species (including empirical evaluation of sequence similarity scores to distinguish real from spurious hits), which in itself provides evidence for the existence of predicted genes through homology. To provide additional evidence for the existence of the novel predicted genes they were subjected to analysis using publicly available RNAseq data from the Sequence Read Archive (SRA)[13]. The datasets used for this analysis are listed in Tables 7 and 8. The tables also show the percentage of the novel genes found that had evidence for their existence in the RNAseq data. For most genomes, most genes predicted by OrthoFiller are supported by RNAseq evidence, with the average percentage of evidence-supported novel genes being 85.3% across the fungal species, and 55.5% across the plant species. Given that the plant RNAseq datasets come from single tissue samples under a single condition it is not expected that all genes will be detected in these samples. For example, similar detection statistics were obtained for the original predicted genes from the source datasets, shown in Tables 7 and 8. It should also be noted that genes that are present in RNAseq reads are more likely to have been annotated already, given that many genome annotation pipelines rely on such data to perform their analyses [3].

**Table 7:** SRA RNA-seq data coverage for novel genes in fungal genomes.

| Species | SRA ID | Instrument/details | Genes in original annotation |  |  | Novel genes |  |  |
| --- | --- | --- | --- | --- | --- | --- | --- | --- |
|  |  |  | Total | W/ reads | % | Total | W/ reads | % |
| *E. gossypii* | N/A<sup>1</sup> | N/A | N/A | N/A | N/A | N/A | N/A | N/A |
| *D. hansenii* | SRR899423 | Helicos Heliscope, single | 5106 | 6739 | 75.77 | 13 | 7 | 53.8 |
| *K. lactis* | SRR1200528 | Illumina Genome Analyzer II, single | 5248 | 5251 | 99.9 | 6 | 6 | 100 |
| *S. cerevisiae* | SRR539284 | Illumina HiSeq 2000, paired end | 6498 | 6572 | 98.87 | 2 | 2 | 100 |
| *Y. lipolytica* | SRR868669 | Illumina HiSeq 2000, single | 7199 | 7520 | 95.73 | 8 | 7 | 87.5 |

<sup>1</sup>No publically available data found for this species

**Table 8:** SRA RNA-seq data coverage for novel genes in plant genomes.

| Species | SRA ID | Instrument/details | Genes in original annotation |  |  | Novel genes |  |  |
| --- | --- | --- | --- | --- | --- | --- | --- | --- |
|  |  |  | Total | W/ reads | % | Total | W/ reads | % |
| *A. thaliana* | SRR3932355 | Illumina HiSeq 2500, paired end.<br>Wild type Columbia rep1 | 162699 | 197160 | 82.52 | 116 | 54 | 46.6 |
| *B. rapa* | SRR2984945 | Illumina HiSeq 2000, paired end.<br>ga-deficient dwarf (gad1-2) +GA<br>rep2 | 180358 | 218457 | 83.56 | 10 | 3 | 30.0 |
| *C. papaya* | SRR3509576 | Illumina HiSeq 2500, paired end.<br>SunUp/Sunset cultivar, young<br>hermaphrodite leaf | 93345 | 112604 | 82.90 | 382 | 309 | 80.1 |
| *C. rubella* | SRR797557 | Illumina Genome Analyzer IIx,<br>paired end | 135008 | 148564 | 90.88 | 228 | 154 | 67.5 |
| *T. cacao* | SRR3217315 | Illumina HiSeq 2000, paired end.<br>Flower/leaf sample | 209407 | 264870 | 79.06 | 94 | 50 | 53.2 |

## Discussion

Here we present OrthoFiller, an automated method for improving the completeness of genome annotations. It leverages information from multiple taxa, clustering genes into orthogroups and finding genes that are conserved between species but that have escaped detection. OrthoFiller is designed to be stringent, conservatively identifying genes that can be confidently identified as missing members of existing orthogroups. Specifically, to pass the filtration criteria for detection by OrthoFiller, genes must be members of orthogroups conserved in multiple species. Thus OrthoFiller will not find genes that lack homologues in other species. These stringent criteria mean that not all genes that could be detected will be detected by the algorithm, but rather that the user should have confidence in the validity of genes identified by the method.

OrthoFiller is intended to be run after a genome annotation is considered by the user to be complete or near-complete. OrthoFiller is designed with small scale genome sequencing projects in mind and is provided to enable users without significant resources for comprehensive RNAseq-based genome annotation to leverage information from related species to improve their genome annotations. However, OrthoFiller is equally suited for use in large-scale genome comparisons, reliably filling gaps in gene sets prior to large scale comparative genomics investigations. Application of OrthoFiller in these cases will enable genes to be analysed in downstream analysis that would otherwise have been classified as absent.

The utility of OrthoFiller is demonstrated on both plant and fungal genome datasets, both in its ability to successfully find missing genes, and in the effectiveness of its filters in eliminating low-quality gene predictions. Application of this method to small groups of plant and fungal genomes resulted in the identification of 570 and 31 genes respectively. These genes are conserved in one or more species but were absent from the genome annotation in which they were predicted. We anticipate that application of OrthoFiller to larger datasets will likely result in further genome annotation improvement. The quality of genes found by OrthoFiller was assessed by artificial removal and recovery of subsets of genes from a single genome, treating those original gene models as true, and evaluating the quality of those genes that were recovered by comparison to the removed genes. In the absence of the OrthoFiller filtration steps, the proportion of poor-quality genes that are recovered is considerably higher.

OrthoFiller is mainly designed for use on genomes that have already undergone some basic level of annotation. As can be seen by comparing the 10% and 90% removal cases in the two data sets, application to very poorly annotated genomes can result in more genes of dubious quality, both from a sequence and an orthogroup perspective. It is worth noting that many of the genes with lower-quality scores, particularly those with only one of the scores being low, can be explained by alternate gene models (in the protein F-score) and shifting of orthogroups due to expansion of proteome sets (in the orthogroup F-score case). In all cases, in the absence of OrthoFiller filtration considerably higher numbers of genes were predicted that didn’t resemble the genes that they were supposed to, indicating that they are erroneous.

The OrthoFiller algorithm is designed to run on a Unix system with python and a minimal number of standard additional tools (HMMer, BedTools, Augustus, R). The speed of the algorithm is principally dependent on the speed of Augustus and HMMer, however processing time can be decreased by parallelising these steps of the method over multiple CPUs.

Accurate and complete genome annotation is of paramount importance to the effective analysis of genomic and transcriptomic data, as well as for phylogenetic inference from genomic data. As the quantity of published genomes increases, care must be taken to ensure accuracy and quality of genome annotations are maintained. Automated methods that leverage publicly available information from multiple species to improve the annotation of newly sequenced genomes will help improve the accuracy and completeness of these resources and thus the quality of all analyses that utilise them.

## Methods

### Data sources

For algorithm development and evaluation, a set of five small, well-annotated fungal genomes (Table 1) and a set of five well-annotated plant genomes (Table 2) were selected. Evaluation of the algorithm focussed on *S. cerevisiae* and *A. thaliana*, as the gene models in these genomes have historically been subject to extensive improvement and revision and are the most likely to be correct.

### Algorithm overview

OrthoFiller proceeds in five stages summarised in Figure 1 and described in detail in the following sections. In brief, the algorithm begins by inferring a set of orthogroups from the protein coding genes of the set of species submitted to OrthoFiller (Figure 1A). The protein sequences in these orthogroups are subject to multiple sequence alignment, converted to nucleotide sequences and used to build HMMs. These HMMs are used to search the genomes of each species under consideration (Figure 1B) and the resultant HMM hits are subject to stringent filtering (Figure 1C) before being used as hints for gene model construction (Figure 1D). The gene models are subject to additional filtering (Figure 1E) and only those gene models that pass all filters are added to the revised genome annotation. The revised genome annotations are then subject to orthogroup inference (Figure 1F) and resultant orthogroups are analysed to confirm the identity of the newly predicted genes. The complete details for each step of this algorithm are described in the sections below.

### Inference of Orthogroups and construction of HMMs

Orthogroups are inferred using OrthoFinder [7]. If a gene from the source annotation is not included in an orthogroup with at least one other sequence, it is classed as a *singleton*, and is not considered in downstream analyses. This is consistent with the problem definition of OrthoFiller, that is to identify unannotated genes that are conserved between species. Amino acid sequences from the orthogroups are aligned with MAFFT [14], using the L-INSI algorithm, and the resultant multiple sequence alignments are back-translated using the source nucleotide sequences. The resulting nucleotide alignments are converted to Hidden Markov models (HMMs) using HMMer [15], each of which is then searched against each input genome in turn to generate a set of hits per HMM per species.

### Evaluation of HMM search results

Due to the probabilistic nature of HMM searches, there is considerable variation in the quality of the relationship between a hit region and the set of sequences used to generate the source HMM. One expects a large amount of “background noise”, that is sequence regions which pass the thresholds of the HMM but whose relevance is dubious. Each HMM hit has an associated bit score, an aggregated base-by-base similarity score between the hit and the aligned sequences used to generate it: we use this score to assess the quality of the hit. The bit score is strongly dependent on the hit length, thus to prevent gene length from biasing downstream analyses the bit score of a hit is divided by the hit length, to generate the *adjusted score* for a hit *h:*

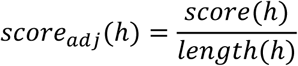

The adjusted score is related to the e-value. However, the e-value calculation enforces a strict lower limit of 1 × 10^−200^, all lower scores being rounded down to zero. Thus use of e-values would introduce irreversible length bias and would lead to downstream errors, as has been shown previously [7]. As bit scores do not have a threshold value, and they have been previously shown to be capable of facilitating accurate inference of phylogenetic trees [16], and length-corrected bit scores are used as the basis of the scoring scheme in OrthoFinder [7], they were used here.

For each species, a threshold value for hit acceptance or rejection based on a hit's adjusted score is created, by considering the distribution of hits which overlapped known genes. Anything above this threshold is considered to be genuine, and anything below this threshold is considered to be noise. An HMM hit is classed as *good* if it overlaps any gene from the orthogroup used to create the HMM, *bad* if it only overlaps genes from orthogroups other than the one used to create the HMM, and *candidate* if it overlaps no known gene at all. Here candidate hits are the potential new genes of interest, and the *good* and *bad* genes are used to inform our judgement about the reliability of the candidate hits.

Distributions of adjusted scores for good and bad hits to the *S. cerevisiae* genome from all HMMs generated by the species in Table 1 are shown in Supplemental Figure 1. Distributions for good and bad hits are clearly delineated into two distinct distributions. Note that in this case there are relatively few candidate hits, since the genome under inspection is already well annotated and is expected to have few missing gene predictions. Skew-t distributions are fit separately to the good and bad score distributions using *gamlss* [17]. Skew distributions were chosen because they allow flexibility in location, shape and scale of the underlying data and are commonly used for estimating parameters such as location and scale, while allowing the same distribution type to be used to fit both the good and bad hits. A separate skew-t distribution for the good and bad hits is fit for each species. In the event that there are insufficient good and bad hits to fit distributions, good and bad hits from the other species are aggregated and a threshold value is calculated from this.

For a given adjusted score *x*, the distributions of the *good* and *bad* hits are used to estimate both the absolute probabilities of a hit being genuine or being a mistake. We can estimate

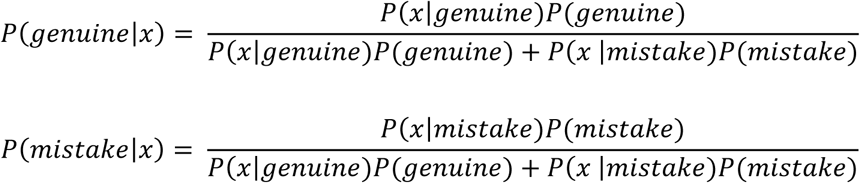

and then retain the hit depending on whether it has a higher probability of being genuine that being a mistake, based on its adjusted score. The probabilities *P*(*genuine*) and *P*(*false*) are estimated by considering the proportion of good/bad hits which are good and bad respectively. The probability density functions *P*(*x*|*genuine*) and *P*(*x*|*false*) are determined using the fitted distributions as described above.

### Acquisition and evaluation of putative predicted genes

Hits which survive the hit filtration step are passed to the gene-finding program Augustus as *hints* specified as exon parts. Only predicted genes that have a nonzero overlap with these hints are retained. These predicted genes are then subjected to a *hint filter*, which aims to separate those genes which have genuinely arisen from the hint from those that overlap the hint by chance. The hint filter evaluates a *hint F-score* for each predicted gene, by comparing against the hints from a particular orthogroup which overlap it. The hint F-score is a measure of how well the found gene corresponds to the hints used to inform its discovery. Each predicted gene *G* will have at least one *hint region* corresponding to it, which is a set of non-overlapping coordinates obtained from merging all hints that overlap *G*, and which are all derived from the same orthogroup. For a hint region *H* and a predicted gene *G*, the hint F score is defined as:

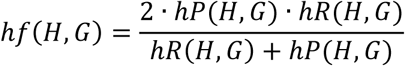

where

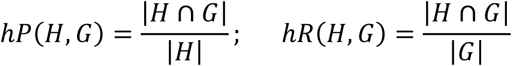

The filter uses a threshold hint F-score value of 0.8 (i.e. on average 80% of the length of the predicted gene is covered by the hit and vice versa), below which potential gene models are discarded. This value was chosen based on an analysis of hint F-scores of good and bad hits (as defined above) versus the Augustus output corresponding to them. Distributions for hint F-scores for the good and bad hits can be seen in Supplemental Figure 4, in which it can be clearly seen that practically all genuine hints pass the threshold value of 0.8.

Once gene models have been filtered, they are fed once again into OrthoFinder, to cluster them into orthogroups. The orthogroup of each newly predicted gene is compared with the orthogroup(s) which were used to predict that gene. It is possible that multiple orthogroups informed the prediction of the same gene; similarly, there may be fluctuations in orthogroup membership between the original and new genomes. It is therefore only required that the new orthogroup into which the gene is clustered has non-zero overlap with at least one of the orthogroups used to predict it, and genes which do not fulfil this criterion are discarded.

### Algorithm evaluation

#### Recovery of removed genes

The test set of species from Table 1 was used to analyse the effectiveness of OrthoFiller for genomes of various levels of completion. Altered versions of the *S. cerevisiae* genome annotation were constructed with 10% and 90% of genes randomly removed, and the level of recovery of the removed genes upon implementation of OrthoFiller was assessed, where a gene ***G*** was considered to be *recovered* if OrthoFiller predicted a gene ***G′*** such that ***G*** and ***G′*** have non-zero overlap.

The quality of the predicted genes was assessed by considering two scores: the orthogroup F-score and the protein F-score. The protein F-score is defined as

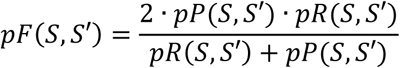

where *S* is the original amino acid sequence and *S′* is the amino acid sequence of the recovered gene, and

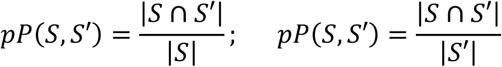

where the intersection is defined to be the sum of identical amino acids in an alignment (MAFFT L- INSI) of the two sequences. The orthogroup F-score is defined as

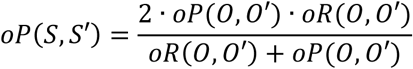

where *0* is the orthogroup that the gene is placed when no deductions have been made, *0′* is the orthogroup into which the gene is placed when OrthoFinder is run on the OrthoFiller results, and

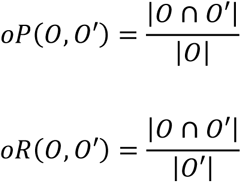

where cardinality of the orthogroups takes into account only genes which were present in the input set of genome annotations, i.e. not counting the newly discovered genes.

### Evaluation of novel predicted genes

RNA-seq data was downloaded from the Sequence Read Archive, and aligned to the genome with BowTie2 using default parameters. Coverage was calculated using BedTools coverage.

### Availability of data and materials

The software is available under the GPLv3 licence at https://github.com/mpdunne/orthofiller.

## Acknowledgements

NA.

## Author's Contributions

SK conceived the project. MPD developed the algorithm. SK and MPD analysed the data and wrote the manuscript. Both authors read and approved the final manuscript.

## Supplemental Figure Legends

**Supplemental Figure 1:**
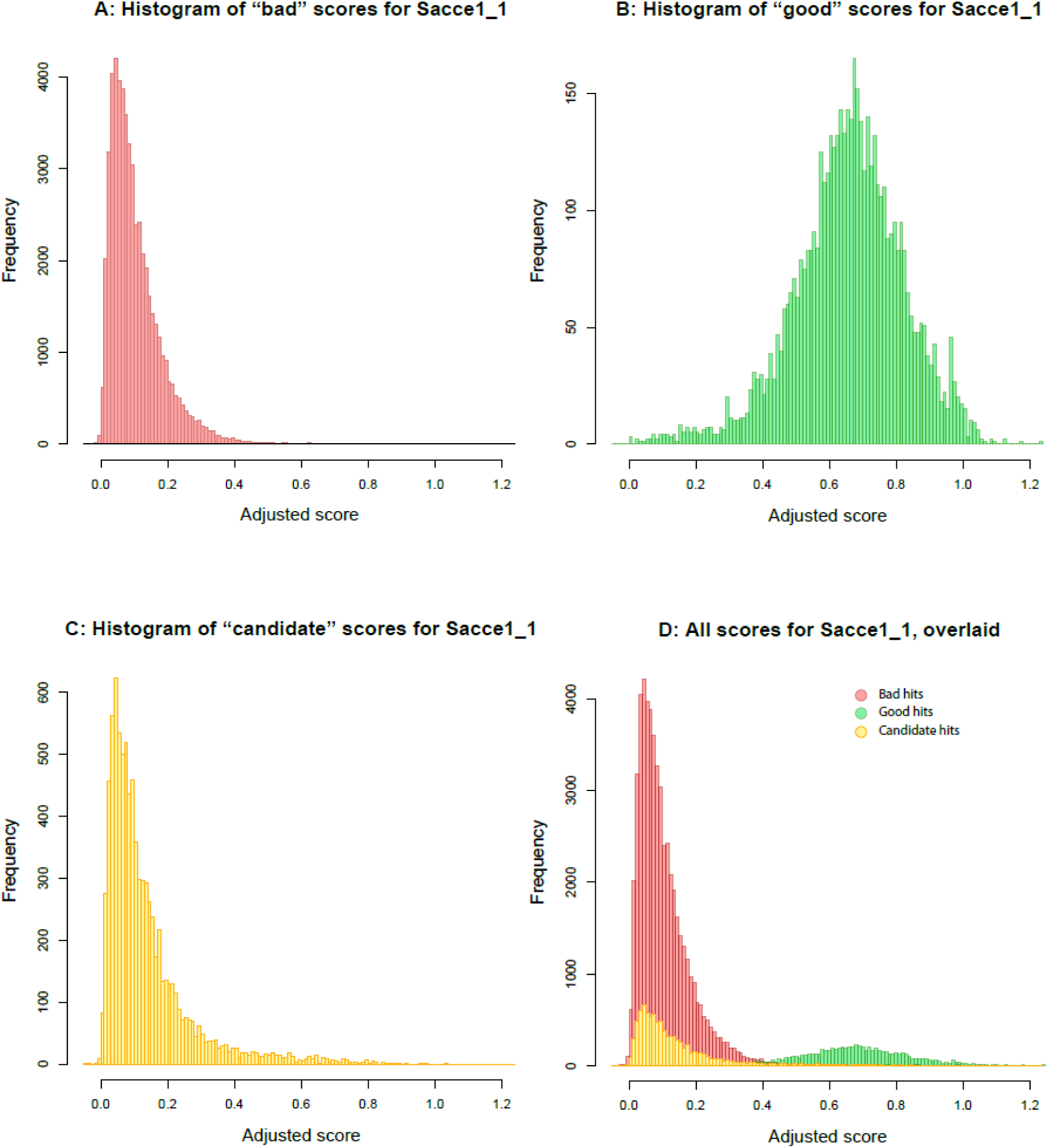
hit score distributions for *good, bad* and *candidate* hits. Hits are to the S. *cerevisiae* genome, using HMMs from all orthogroups. **A)** Length normalised bit scores of HMM hits to regions of the genome that contained genes that were not part of the orthogroup used to generate the HMM (bad hits). **B)** Length normalised bit scores of HMM hits to regions of the genome that do contain the gene used to generate the HMM (good hits). **C)** Length normalised bit scores of HMM hits to regions of the genome that do not contain any previously annotated genes (candidate novel gene hits). **D)** All distributions overlaid.

**Supplemental Figure 2:**
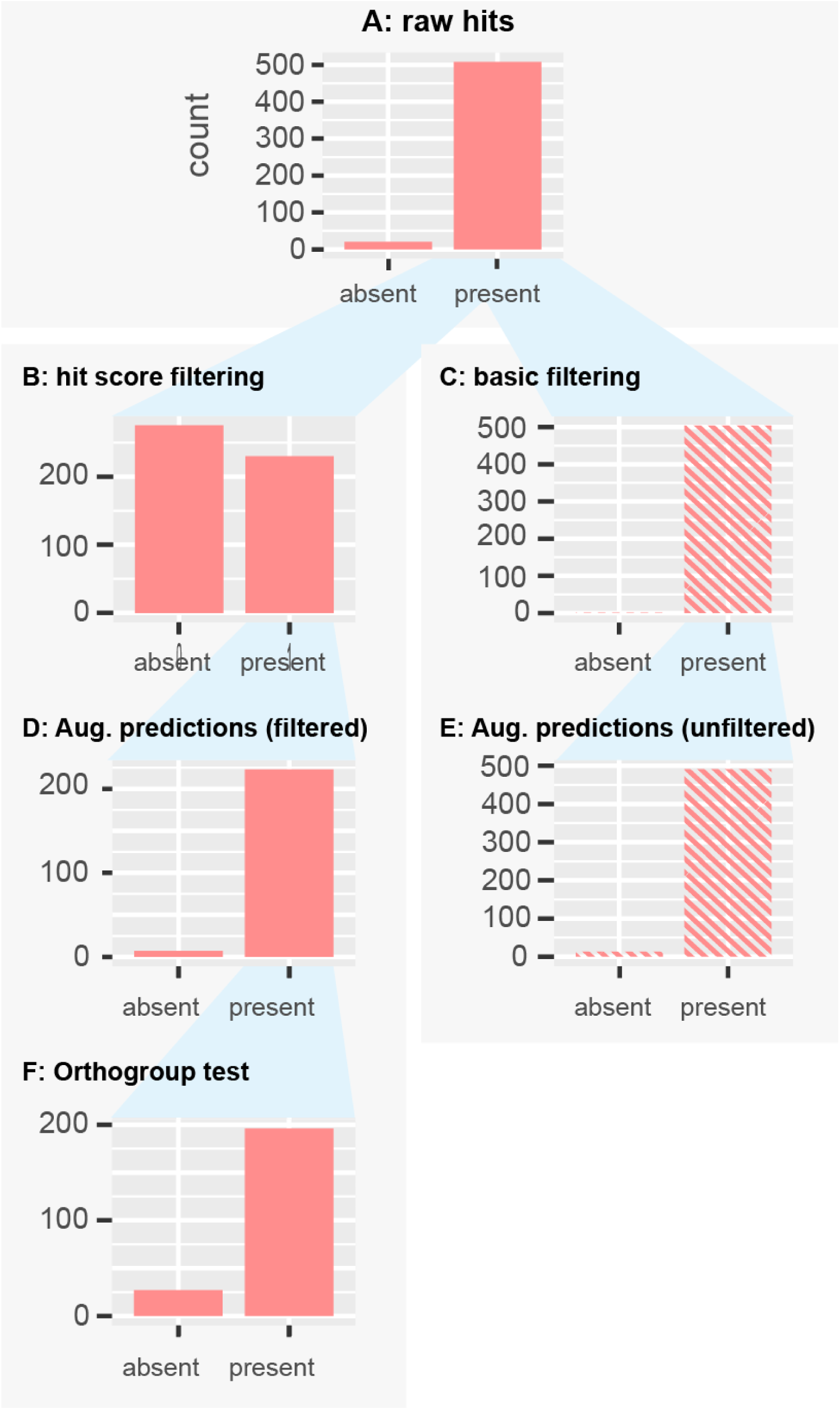
Recovery of removed genes from *S. cerevisiae* after 10% removal: Representation of removed genes at each stage, filtered vs. unfiltered cases. **A)** The number of deleted genes that obtained hits from one or more orthogroup HMMs. **B)** The number of deleted genes that had hits after OrthoFiller hint filtration. **C)** No hint filtration. **D)** The number of deleted genes for which a gene prediction was made using Augustus that satisfied OrthoFiller filtration tests. **E)** The number of deleted genes that for which a gene prediction was made using Augustus in the absence of OrthoFiller filtration. **F)** The number newly predicted genes that were retained or discarded based on the orthogroup assignment filter step in OrthoFiller.

**Supplemental Figure 3:**
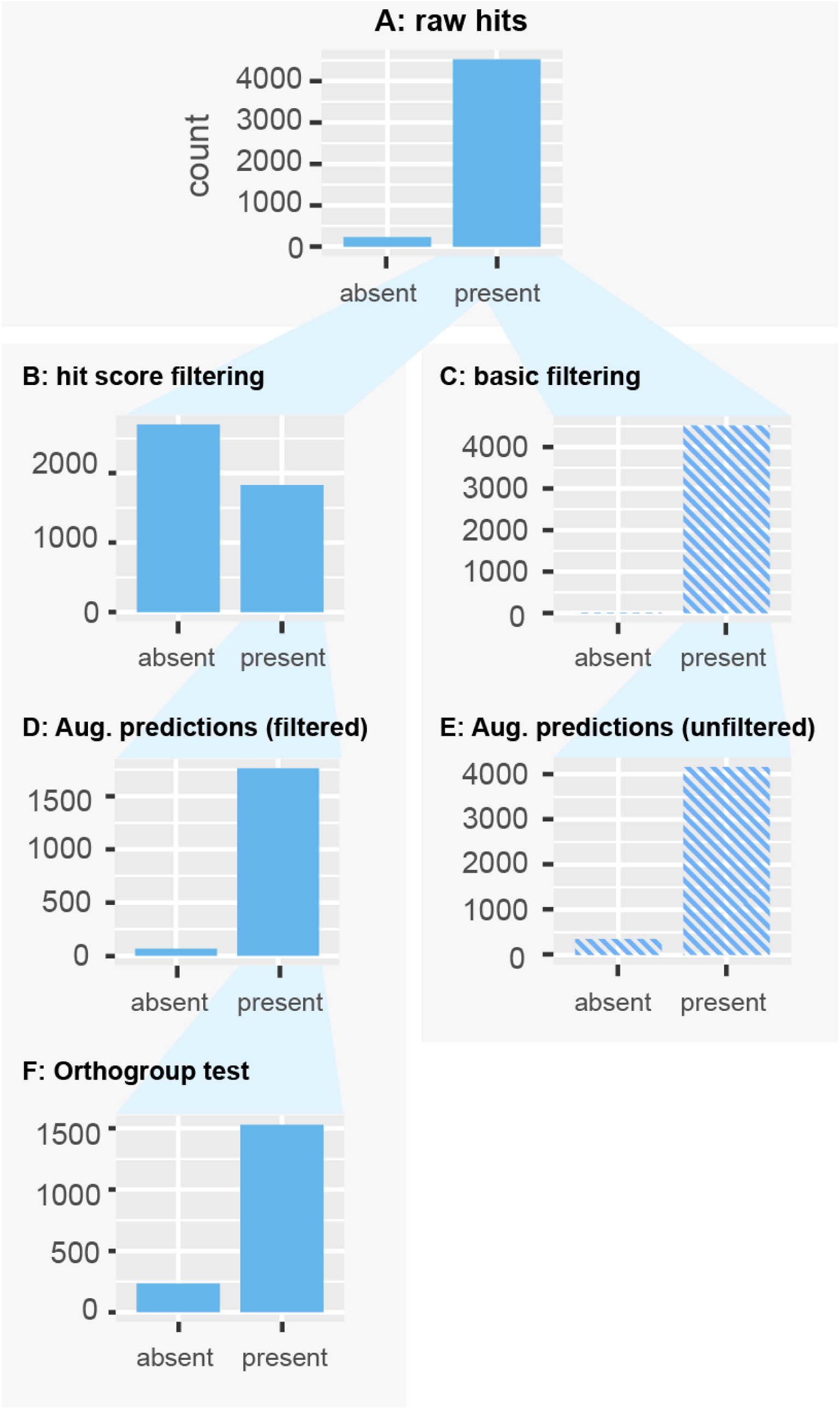
Recovery of removed genes from *S. cerevisiae* after 90% removal: Representation of removed genes at each stage, filtered vs. unfiltered cases. **A)** The number of deleted genes that obtained hits from one or more orthogroup HMMs. **B)** The number of deleted genes that had hits after OrthoFiller hint filtration. **C)** No hint filtration. **D)** The number of deleted genes for which a gene prediction was made using Augustus that satisfied OrthoFiller filtration tests. **E)** The number of deleted genes that for which a gene prediction was made using Augustus in the absence of OrthoFiller filtration. **F)** The number newly predicted genes that were retained or discarded based on the orthogroup assignment filter step in OrthoFiller.

**Supplemental Figure 4:**
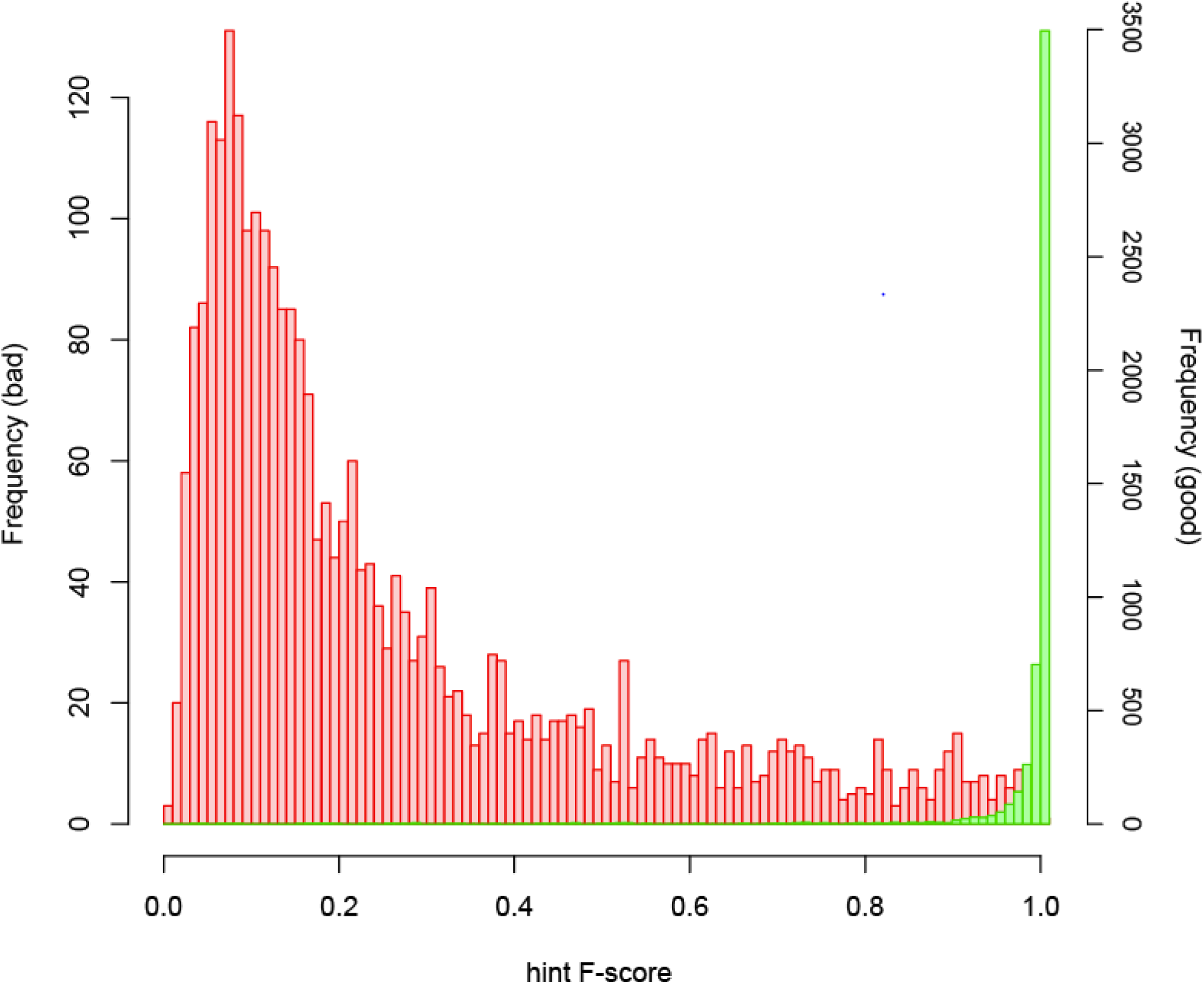
Distribution of hint F-scores for good vs. bad hints. Here, Augustus has been allowed to predict genes that are already present in the input genome, hence we can consider separately the good and bad hits as hints. Shown are the distributions of hint F-scores for good (green) and bad (red) hits respectively, demonstrating that practically all of the genuine hints have a hint F-score of 0.8 or higher.

